# Assessment of Environmental DNA Surveys for the Cryptic Salamander Mussel (*Simpsonaias ambigua*)

**DOI:** 10.64898/2026.02.10.705175

**Authors:** Nate Marshall, Noah Berg, Triston Mullins, Christine Stahlman, Cheryl Dean, Mae Sierra, Cody Fleece

## Abstract

*Simpsonaias ambigua* (Salamander Mussel), is a small and thin shelled freshwater mussel often found in difficult to survey habitats, such as beneath slab stones, in the crevices of rock walls, or buried within roots of emergent vegetation and in undercuts of banks. The use of environmental DNA (eDNA - genetic material released from urine, waste, mucus, or sloughed cells) sampling may improve detection and assessment of presence / absence for this rare mussel in comparison to visual tactile techniques. This study completed side by side comparisons of traditional mussel searches and eDNA for a direct assessment of mussel detection efficiencies. Surveying was conducted in several waterbodies of different habitat characteristics with varying abundances of *S. ambigua*. Additionally, a broad assessment of *S. ambigua* presence was conducted throughout the proposed critical habitat reach of the Blanchard River in northwest Ohio, to assess if the species remained extant. All eDNA samples were also assessed for the presence of *Necturus maculosus* (Mudpuppy), the obligate host species for *S. ambigua*. The eDNA sampling successfully detected *S. ambigua* from multiple sites and watersheds where it was found with visual surveys. In some cases, eDNA detections occurred in locations where over 16 hours of search yielded only a single individual or fresh dead shells, supporting the sensitivity of eDNA for detection of rare species. Furthermore, probability of detection analysis suggests eDNA sampling can provide high detection efficiency with relatively low effort in comparison to visual searches. The development and validation of an eDNA protocol for the simultaneous detection of *S. ambigua* and its host salamander increases survey efficiency, reduces field costs, and can support future conservation efforts for listing drainages of extant populations and monitoring conservation goals.

## Introduction

Of the 303 currently recognized mussel species within the United States (US) (FMCS 2023), nearly one third are federally endangered (77 species) or federally threatened (18 species) (USFWS 2023). Due to their rarity and the fact that unionids burrow into the benthic substrate, traditional visual surveys are time-consuming and require expert taxonomists. Therefore, novel methods to improve the efficiency and accuracy of surveys could be beneficial for population management.

The analysis of environmental DNA (eDNA - genetic material released from urine, waste, mucus, or sloughed cells) is increasingly integrated into natural resource surveys designed to detect the presence of special status species and describe entire assemblages (Beng & Corlett 2020, Deiner et al. 2021).Environmental DNA surveys can be more sensitive, less costly, less intrusive on the environment, and provide improved sampling capabilities for challenging and remote habitats (Jerde 2021, Sternhagen et al. 2024). Therefore, eDNA offers great potential as an efficient method to assess mussel assemblage in river systems (Marshall et al. 2022, 2026, Nogueira et al. 2025).

There are several different molecular methodologies for analyzing eDNA samples, each with varying costs and inherent limitations. The various laboratory methodologies are often generalized into two categories: (1) targeted approaches that utilize species-specific assays to detect a single species of interest and (2) broad community characterization approaches that utilize conserved genetic markers to simultaneously detect entire community assemblages (Bylemans et al. 2019, McColl-Gausden et al.2023). The most common species-specific targeted approaches include quantitative PCR (qPCR) (Langlois et al. 2021), digital droplet PCR (ddPCR) (Baudry et al. 2023), and most recently targeted high-throughput sequencing (McCarthy et al. 2023). Broad community characterization approaches typically refer to *metabarcoding* which require high-throughput sequencing technologies (Ruppert et al. 2019). It is often suggested species-specific assays are more sensitive for the detection of rare DNA targets (Wood et al. 2019, Blackman et al. 2020).

*Simpsonaias ambigua* (Salamander Mussel), is a small (approximately 50 mm length), thin shelled mussel. This species is typically found in difficult to survey habitats, such as beneath large, flat stones or in the crevices of rock walls, as well as among the roots of emergent vegetation, large woody debris, and undercuts of banks (Bogan 2009, Porto-Hannes et al. 2025). Additionally, it is unique among freshwater mussels, as it is the lone unionid to utilize an amphibian as a host species, *Necturus maculosus* (Mudpuppy) (Porto-Hannes et al. 2025). Its utilization of crevices and undercut bank habitat is likely tied to the habitat preference of its host amphibian, but *S. ambigua* has also shown a preference for darkened shelter and to aggregate in groups (Stegman 2000).

In the US, *S. ambigua* has been proposed endangered as 98.5% of populations have been deemed at risk of extirpation (USFWS 2023). Additionally, 37 river units in the US have been proposed as critical habitat for *S. ambigua* (USFWS 2023), which is defined by the United States Fish and Wildlife Service (USFWS) as ‘*specific areas occupied by the species at the time it was listed, that contain the physical or biological features that are essential to the conservation of endangered and threatened species and that may need special management or protection*’ (USFWS 2017 - fws.gov/sites/default/files/documents/critical-habitat-fact-sheet.pdf). The development and implementation of eDNA methodology for detecting this species and its obligate host salamander will likely aid in obtaining a better understanding of its realized distribution and provide improved presence / probable absence assessments.

Previous studies have demonstrated successful eDNA collection and detection for this species when using qPCR analysis (Porto-Hannes et al. 2023), which has included the development of an eDNA protocol in Canada to complement visual surveys (Porto-Hannes et al. 2021) and eDNA detections leading to new visual observations (Douglass et al. 2025). However, the laboratory analyses employed here included both metabarcoding, to document complete mussel assemblage, as well as qPCR, to assess detection sensitivity. Surveying occurred at sites in five of the proposed critical habitat units (Figure 1, Table 1), which included sites of varying habitat type (e.g., crevices of rock walls vs undercut banks) and ranges of mussel abundance. Sampling occurred at 13 locations throughout the designated critical habitat within the Blanchard River, OH, at the location of the only known record of occurrence in that water body (USFWS 2023). The eDNA samples were also analyzed for the detection of *N. maculosus* to further assess the potential for sites to support a population of *S. ambigua*.

**Figure 1.**
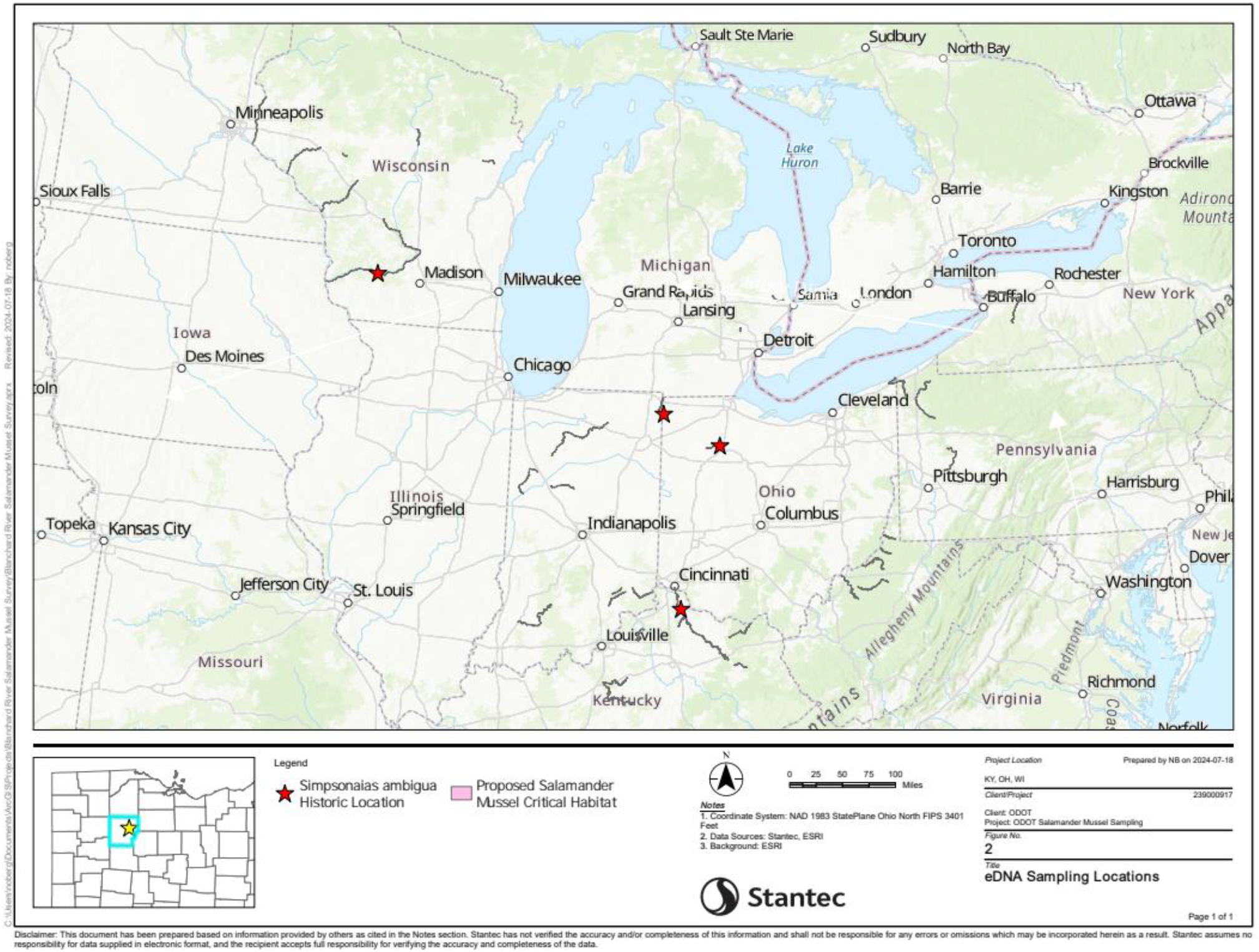
Sampling locations for visual and eDNA surveys for Simsponaias ambigua and Necturus maculosus within the Wisconsin River, South Fork Licking River, Licking River, Fish Creek, and Blanchard River.

**Table 1.** Number of environmental DNA (eDNA) and visual surveys conducted in each watershed for assessment of Simsponaias ambigua.

| **Watershed** | **Critical Habitat**<br>**Unit No.** | **No. eDNA Sites** | **No. Visual Sites** | **Sampling Dates** |
| --- | --- | --- | --- | --- |
| Blanchard R. | 13 | 14 | 3 | 08/22/2024-08/24/2024 |
| Fish Cr. | 12 | 3 | 2 | 08/26/2024-08/27/2024 |
| Licking R. | 27 | 1 | 0* | 08/19/2024 |
| South Fork Licking R. | 28 | 3 | 2 | 08/19/2024-08/20/2024 |
| Wisconsin R. | 8 | 1 | 0* | 08/02/2024 |

*\*Simsponaias ambigua* presence known from previous survey data

## Methods

### eDNA Sample Collection

Environmental DNA sampling was conducted within five watersheds, which included the Blanchard River and Fish Creek in Ohio, Licking River and South Fork Licking in Kentucky, and the Wisconsin River in Wisconsin (Table 1 & Figure 1). The number of survey sites varied between watersheds ranging from one to 14, with the highest occurring within the Blanchard River (Table 1). Concurrent visual tactile surveys were conducted at seven sites.

At sites without a visual mussel survey, anywhere from three to ten replicate water samples were collected (see Appendix A). Water samples were collected from near the benthos using a peristaltic pump with a polypropylene filter holder attached to a painter’s pole. A new polypropylene filter holder was used between all samples collected on each day. eDNA was collected along a transect spanning the width of the river for each replicate sample. Once the first sample was collected, the surveyor took the next sample ∼2-3m upstream, and continued this for three replicates per site. Samples were filtered on a 47-mm-diameter GF/C. Each sample consisted of filtering 1000mL of river water. Filters were placed into separate coin envelopes, which were then placed in Ziploc bags with silicone desiccant beads and stored in a freezer (−20^°^ C).

At sites with a corresponding visual mussel survey, four replicate water samples were collected. These samples were collected as described above, however each replicate was spaced every 10 m to align with the search cells of the visual survey (see below).

On each day of sampling, at least one negative field control was collected consisting of 500mL of distilled water processed along with field samples. Sampling was always conducted in the upstream direction, to avoid potential contamination and sediment disturbance. Field staff used new gloves between each sample, and all equipment (e.g., polypropylene filter holders, extendable pole, peristaltic pump tubing) was washed with 30% bleach following each day of field collection. All filters were shipped on ice to Cramer Fish Science Genidaqs (Sacramento, California, USA, https://genidaqs.com) for DNA extraction and analysis.

### Visual Surveys

A visual tactile survey was conducted at a subset of eDNA survey sites. This subset of sites was determined based on the presence of suitable *S. ambigua* habitat during the time of eDNA sampling, with any visual survey occurring within 1000m of a bridge or public access point. Each visual tactile survey consisted of eight search cells, with each extending 10m long and spanning half the width of the river, for a search cell of approximately 100m^2^. Within each search cell, three 20-minute passes occurred, for a total search effort of one hour per search cell and eight hours per site. Field personnel searched for mussels using a Self-Contained Underwater Breathing Apparatus (SCUBA) or a snorkel. Searches included moving cobble and woody debris, tactile exploration of undercut banks or rock crevices, hand sweeping away silt, sand, and/or small detritus, and disturbing/probing the upper five centimeters (cm) of substrate. During each search mussels were collected in mesh bags and brought to shore for identification and data collection. Mussels were identified to the species level, length was measured to the nearest millimeter (mm), and individuals were sexed (if sexually dimorphic). After all passes were completed within a search cell, all mussels were returned to the approximate location where found. Visual tactile searches were not conducted at the Wisconsin River site as part of this study, but prior work had documented a very large extant population of *S. ambigua* (Stantec 2022, 2024).

### Sample DNA Extraction and Inhibition Removal

DNA was extracted from each filter using a Qiagen DNeasy® Blood and Tissue Kit following manufacture’s protocol except as documented below. Each filter was incubated overnight at 56 °C in 540 µl Buffer ATL and 60 µl Proteinase K. The resulting supernatant was passed through a Qiashredder spin column, then mixed with 600 µl Buffer AL and incubated at 70 ºC for 10 min. After adding 600 µl 100% ethanol, the resulting mixture was loaded onto a DNeasy Spin column. DNA wash stepsfollowed manufacture’s protocol. The final elution volume was 100 µl. A negative control was simultaneously extracted to test for possible laboratory contamination.

A potential issue with eDNA analysis arises when inhibitor compounds are present within a sample from humic, phytic, and tannic acids, leaf litter, algae, and sediments. These compounds reduce the efficiency and accuracy of the PCR (Lance & Guan 2020). High levels of total suspended solids and high concentrations of disturbed sediment may reduce eDNA quantification and increase presence of PCR inhibitors (Stoeckle et al. 2017). The DNA was further processed with a Zymo Research One Step PCR Inhibitor Removal kit (Zymo Research, Irvine, California, USA) to mitigate potential inhibition.

Samples collected from watersheds that failed to obtain a detection of *S. ambigua* with qPCR were further assessed for inhibition with a spike-in trial using a known quantity of *Hemiramphus brasiliensis* (Ballyhoo), an Atlantic Ocean bait fish that is absent from any of the sampling watersheds. In brief, 1.4 µl of *H. brasiliensis* DNA (0.002 ng/µl) was added to 12.6 µl of field sample, or DNA-free water (control), Spiked samples and water controls were used as template in a qPCR reaction for *H. brasiliensis*, using an assay targeting the cytochrome b gene region (Bowen et al. 2024). Reactions were performed in triplicate 10µl reactions containing 4 µl spiked template, final concentration 900nm forward and reverse primers, 250nm probe, and 1X Applied Biosystems™ TaqMan™ Environmental Master Mix 2.0. The amplifications started with an initial denaturation at 95°C for 10 min, followed by 40 cycles of 95°C for 15s and 60°C for 60s.Samples were analyzed for inhibition by comparing Cq values of spiked samples with the average Cq value of spiked DNA-free water. Samples were determined to not be inhibited, as no samples demonstrated a ΔCq (CqSpiked water – CqSpiked sample) ≥3, the point at which inhibition is generally accepted to be present (Hartman et al. 2005).

### Metabarcoding

For metabarcoding community analysis, each water sample was amplified for a ∼175 base pair (bp) fragment of the mitochondrial 16S gene region using an assay developed by Prié et al. (2021) and previously validated for detection of freshwater mussels from eDNA samples (Marshall et al. 2022).

Library preparation followed a three-step PCR process originally described in Marshall et al. (2022) with changes to the initial PCR step to improve success in environments with PCR inhibitors. An initial library preparation using Qiagen Plus Multiplex Master Mix failed to produce usable data, but a new preparation using Applied Biosystems™ TaqMan™ Environmental Master Mix 2.0 met all quality control measure.This is the same master mix as used in qPCR inhibition testing and appears to increase PCR success for this study. Additionally, the initial PCR total volume was increased to 20ul from suggested 10ul to mitigate against potential inhibitors (dilution effect).

Initial PCR amplification was completed for each sample in triplicate with 20 µl PCR reactions containing 4 µl extracted eDNA, 0.4µM of each primer, and 1X Applied Biosystems™ TaqMan™ Environmental Master Mix 2.0. The amplifications started with an initial denaturation at 95ºC for 5 min, followed by 35 cycles of 95ºC for 15s, 5% ramp down to 55ºC for 30s, and 72ºC for 30s. Triplicate PCR products were diluted 1:10 then pooled prior to starting the Illumina adaptor and barcoding PCR processes.

The MiSeq library dual indexed paired-end sample preparation was adapted as described in Miya et al. (2015) from ‘16S metagenomic sequencing library preparation: preparing 16S ribosomal gene amplicon for the Illumina MiSeq system (Illumina part no. 15044223 Rev. B, San Diego, California, USA). A PCR process initiated the incorporation of Illumina adaptors and multiplexing barcodes using Prié et al. (2021) forward and reverse primers containing 33 or 34 base pairs of 5’ Illumina hanging tails to provide a priming site for a final PCR to incorporate barcodes and remaining base pairs of Illumina adaptors. The 12 µl PCR reaction contained 2 µl diluted pooled PCR product, 0.3 µM Illumina adaptor primers and 6 µl 1X Qiagen Plus Multiplex Master Mix. The PCR process denatured for 95ºC for 5 min, 5 cycles of 98ºC for 20s, 1% ramp down to 65ºC for 15s, and 72ºC for 15s, followed by 7 cycles of 98ºC for 20s, 5% ramp down to 65ºC for 15s, and 72ºC for 15s. PCR product was diluted 1:10 prior to use in the barcode adaptor PCR process.

The final PCR incorporated paired-end dual indices (eight base pair barcodes) that allowed samples to be identified in the raw read data, and the p5/p7 adaptor sequences to allow the sample to bind onto the Illumina MiSeq flow cell. This final 12μl PCR reaction contained 1μl diluted product from the previous PCR, 0.3 µM forward and reverse indexed primer and 6ul 1X KAPA HiFi HotStart Ready Mix PCR Kit (Roche Diagnostics, Indianapolis, Indiana, USA). Conditions were 3 minutes of initial denaturation at 95°C, followed by 10 cycles at 98°C for 20 s, 5% ramp down to 72°C for 15 s, with a final 5 min 72°C extension. All PCRs were completed on Bio-Rad C1000 Touch Thermal Cyclers. Illumina adapted PCR products were pooled with equal volumes, then size selected (target ∼319bp) using 2% agarose gel electrophoresis. The final pool was sequenced with 2× 300 nt V3 Illumina MiSeq chemistry by loading 6.4 pmol library. An additional 20% PhiX DNA spike-in control was added to improve data quality of low base pair diversity samples. Additionally, a PCR no-template negative control was run for each library preparation step.

### Bioinformatic Processing & Taxonomic Identification

The metabarcoding data was processed following a bioinformatic pipeline previously described in Marshall et al. (2022). The MiSeq runs were processed separately. The forward and reverse primer sequences were removed from the demultiplexed sequences using the cutadapt (Martin et al. 2011) plugin within QIIME 2 (Bolyen et al. 2019). Next, sequence reads were filtered and trimmed using the denoising DADA2 (Callahan et al. 2016) plugin within QIIME 2. Based on the quality scores from the forward and reverse read files, a “truncLen” was set to 120 for the forward and 110 for the reverse read files. Using DADA2, error rates were estimated, sequences were merged and dereplicated, and any erroneous or chimeric sequences were removed. Unique sequences were then clustered into Molecular Operational Taxonomic Units (MOTUs) using the QIIME 2 vsearch de-novo with a 97.5% similarity threshold (Coghlan et al. 2021, Marshall et al. 2022). MOTUs from unionid taxa were identified to the species-level using the Basic Local Alignment Search Tool (BLAST+, https://blast.ncbi.nlm.nih.gov/Blast.cgi; Camacho et al. 2009) against our custom database of both in-lab generated sequences and mt-16S sequences downloaded from NCBI GenBank. These MOTUs were further validated with comparisons against the complete NCBI nr database, to investigate alignment to mis-labeled sequences or species not historically within the sampling region.

Freshwater mussels display a unique form of mitochondrial inheritance, termed doubly uniparental inheritance (DUI), in which males possess a paternal mitochondrial mitotype that is largely restricted to male gonads and gametes (Gusman et al. 2016). As the male mitotype is genetically distinct from the female mitotype (Curole & Kocher 2005), detections originating from the female or male mitotype were separated within the metabarcoding dataset. The male lineage for *S. ambigua* was able to be detected in the current study because a voucher swab was collected and used to generate the first reference sequence, whereas a majority of species lack reference genetic sequences for their male lineage and cannot currently be detected within an eDNA dataset (Marshall et al. 2025).

Sequences were assigned to a species if they met a threshold of >97.5% identity and 100% query coverage. Furthermore, sequences that were assigned to multiple species with the same BLAST e-value score were inspected and a final decision was made based on known distribution and presence within the sampled watershed. Additionally, if multiple sequences were assigned to the same taxonomy, they were inspected and removed or collapsed into a single MOTU to obtain a final matrix of read counts per taxa.

In metabarcoding analysis, the sequencing process can introduce a form of sample cross-contamination in which sequences from one sample are falsely detected within another sample, often termed ‘critical mistags’, ‘tag jumps’, or ‘index hopping’ (Carlsen et al. 2012; Esling et al. 2015; Bohmann et al. 2022, Richardson 2022). These sequencing artifacts generally represent a small fraction of the total sequences (Esling et al. 2015; Schnell et al. 2015), yet they can lead to false detections in some circumstances.Therefore, the final processing of the MOTUs in this study consisted of estimating and removing mistags from the dataset following the framework outlined by Richardson (2022). Based on the number of reads per MOTU within the field and laboratory negative controls, the mistag rate was estimated for all dual-index combinations. The estimated mistag rate of the current dataset was 0.001, but a conservative estimate of 0.0075 was used for identifying and removing mistags (Richardson 2022). Any sequences identified as field contamination were not included in estimating the mistag rate (see Results).

### qPCR

Samples were processed with qPCR for the detection of *S. ambigua* and *N. maculosus*. The presence of *S. ambigua* DNA was assessed using a previously developed qPCR assay targeting the mt-ND1 gene region (Porto-Hannes et al. 2023). The presence of *N. maculosus* DNA was assessed using a previously developed qPCR assay targeting the mt-COI gene region (Collins et al. 2019, Sutherland et al. 2020).

qPCR amplifications were completed in triplicate 10 µl reactions. *S. ambigua* contained 4 µl extracted eDNA, final concentration of 300nM forward and reverse primers, 200mM probe, and 1X Applied Biosystems™ TaqMan™ Environmental Master Mix 2.0. *N. maculosus* contained 4 µl extracted eDNA, final concentration of 900nM forward and reverse primers, 100mM probe, and 1X Applied Biosystems™ TaqMan™ Environmental Master Mix 2.0. For both assays, the amplifications started with an initial denaturation at 95ºC for 10 min, followed by 40 cycles of 95ºC for 15s and 60ºC for 60s. Three positive controls and three no-template controls were assessed on each qPCR plate. Each eDNA sample was analyzed with three technical replicates with qPCR. Any sample that had no detection with qPCR for *S. ambigua* but was positive for the female mitotype with metabarcoding was re-analyzed with an additional three technical replicates.

### Estimating eDNA Detection Probability

Occupancy estimation is a model-based approach to estimate the probability of species presence in an area while accounting for the imperfect detection probabilities that are inherent in most sampling methods (MacKenzie et al. 2002). Using the eDNA data, single-season occupancy models were evaluated for *S. ambigua* to estimate the mean detection probability (i.e., the probability of successful eDNA detection of a species within a replicate environmental sample). Models were evaluated for both qPCR and metabarcoding data as described in the following sections.

### Metabarcoding

The probability of eDNA detection was evaluated from the metabarcoding data for only the female mitotype of *S. ambigua*. Metabarcoding occupancy models were analyzed in the R package ‘unmarked’ (Fiske & Chandler 2011). Simplified null models were used to assess probabilities of eDNA detection for each species (i.e., occupancy models were analyzed with no covariates added). The occupancy models for metabarcoding included two hierarchical levels within our sampling design, which included (ψ)(*p*):

ψ – the probability of eDNA occurrence at the site-level, and

*p* – the conditional probability of eDNA detection in a sample given that the eDNA is present at the site-level.

The probability of eDNA detection in this framework is dependent on:

(1) the number of sample replicates (x) collected at a site: *p*.x = 1-(1-*p*)^x^

### qPCR

The probability of eDNA detection was evaluated from the qPCR data for both *S. ambigua* and *N. maculosus*. qPCR occupancy models were analyzed in the R package ‘eDNAoccupancy’ (Dorazio & Erickson 2018). Simplified null models were used to assess probabilities of eDNA detection for each species (i.e., occupancy models were analyzed with no covariates added). The occupancy models for qPCR included three hierarchical levels within our sampling design, which included (ψ)(Θ)(*p*):

ψ – the probability of eDNA occurrence at the site-level,

Θ – the conditional probability of eDNA collection in a sample given that the eDNA is present at the site-level, and

*p* – the conditional probability of eDNA detection in a laboratory technical replicate given that the eDNA is present in the sample.

The cumulative probability of eDNA detection (Θ.*p*) with qPCR was defined as the combined probability of collecting eDNA within a sample (Θ) and subsequently detecting that eDNA within a laboratory technical replicate (*p*). This cumulative probability of eDNA detection in this framework is dependent on:

1. the number of technical replicates (x) analyzed per sample: *p*.x = 1-(1-*p*)^x^, and
2. the number of water sample replicates (y) collected at a site: Θ.*p* =1-(1-(Θ\**p*.x))^y^

## Results

### Field Contamination & Mistags

Detection of *S. ambigua* and *N. maculosus* were never recorded in any of the field or laboratory controls. The field negative control collected at the Wisconsin River site had high sequence counts for two species, *Lampsilis cardium* (6236 sequence reads) and *Potamilus fragilis* (2392 sequence reads). The eDNA sampling at this site occurred at the end of the day following a mussel relocation survey, and the eDNA sampling was conducted by the same field staff that conducted the scuba mussel surveys. These two particular species were common during the mussel relocation survey that day. It is likely that the high sequence read counts observed in this field control are the result of residual DNA on the surveyors stemming from the mussel relocation. As these species were observed and handled at the site that day, their detections were included within the field eDNA samples. This is the only site in which mussel surveys occurred prior to eDNA collection and the only field control with high sequence counts (the next highest amount amongst field controls was 437 total mussel sequence counts).

After removing the sequence counts for *L. cardium* and *P. fragilis* from the Wisconsin River field control, the amount of mussel sequence reads per control sample ranged from 0 to 690. In total, 22 MOTUs were detected on 122 occurrences within the 27 field and laboratory controls. These detections were all considered the result of mistags and subsequently assessed following Richardson (2022). After identifying sequence read counts falling within the estimated mistag rate (0.0075), only six of the 122 mistags remained within the control samples. From the environmental samples, a total of 12% (258 of 2171) of the detections were classified as being within the range of probable mistag, and thus they were removed.

### Metabarcoding Bioinformatic Processing

In the final dataset, a total of 16,435,999 reads were classified as a unionid female mitotype sequence, while 11,016,053 were classified as a unionid male mitotype sequence. The amount of sequencing reads per sampling watershed ranged from 4,248,148 in Wisconsin River, 3,815,892 in South Fork Licking, 2,026,377 in Licking River, 3,332,685 in Blanchard River, and 3,986,263 in Fish Creek.

### Environmental DNA Detections

A total of 27 eDNA samples out of the 88 collected were positive for *S. ambigua* with either metabarcoding or qPCR (Table 2 & 3). This included positive detections from eight sampling sites, with at least one site being positive per water body (Table 2). Metabarcoding yielded positive detection of *S. ambigua* female mitotype from 20 of those 27 samples (74%), and of *S. ambigua* male mitotype from 19 of those 27 samples (70%) (Table 2 & 3). qPCR yielded positive detection from 21 of those 27 samples (78%) (Table 2 & 3). Of the 27 samples with positive detections, 17 (63%) yielded positive detection for the female mitotype with metabarcoding and a positive with qPCR (Table 2 & 3). Whereas only 12 of those 27 samples (48%) yielded positive detection for both female and male mitotypes with metabarcoding and a positive with qPCR (Table 2 & 3).

**Table 2.** Environmental DNA detection of Simpsonaias ambigua using metabarcoding and qPCR methodologies and detection of Necturus maculosus using qPCR. Detections are listed as the number of positive samples out of the total collected at a site.

| Watershed | Site ID | *S. ambigua* Metabarcoding Reads |  | Positive qPCR Replicates |  |
| --- | --- | --- | --- | --- | --- |
|  |  | Female | Male | *S. ambigua* | *N. maculosus* |
| Wisconsin R. | WR | 9/10 | 8/10 | 9/10 | 9/10 |
| South Fork Licking R. | SFL1 | 1/4 | 0/4 | 2/4 | 1/4 |
| South Fork Licking R. | SFL2 | 0/4 | 4/4 | 2/4 | 1/4 |
| South Fork Licking R. | SFL3 | 3/4 | 4/4 | 3/4 | 3/4 |
| Licking R. | LR | 4/4 | 3/4 | 4/4 | 1/4 |
| Fish Cr. | FC1 | 1/4 | 0/4 | 0/4 | 0/4 |
| Fish Cr. | FC2 | 0/4 | 0/4 | 0/4 | 0/4 |
| Fish Cr. | FC3 | 1/6 | 0/6 | 1/6 | 1/6 |
| Blanchard R. | BR1 | 1/4 | 0/4 | 0/4 | 0/4 |
| Blanchard R. | BR2 | 0/3 | 0/3 | 0/3 | 0/3 |
| Blanchard R. | BR3 | 0/7 | 0/7 | 0/7 | 1/7 |
| Blanchard R. | BR4 | 0/3 | 0/3 | 0/3 | 0/3 |
| Blanchard R. | BR5 | 0/3 | 0/3 | 0/3 | 0/3 |
| Blanchard R. | BR6 | 0/3 | 0/3 | 0/3 | 0/3 |
| Blanchard R. | BR7 | 0/3 | 0/3 | 0/3 | 1/3 |
| Blanchard R. | BR8 | 0/3 | 0/3 | 0/3 | 0/3 |
| Blanchard R. | BR9 | 0/4 | 0/4 | 0/4 | 0/4 |
| Blanchard R. | BR10 | 0/3 | 0/3 | 0/3 | 0/3 |
| Blanchard R. | BR11 | 0/3 | 0/3 | 0/3 | 0/3 |
| Blanchard R. | BR12 | 0/3 | 0/3 | 0/3 | 0/3 |
| Blanchard R. | BR13 | 0/3 | 0/3 | 0/3 | 0/3 |
| Blanchard R. | BR14 | 0/3 | 0/3 | 0/3 | 0/3 |

A total of 18 eDNA samples out of the 88 collected were positive for *N. maculosus* DNA (Table 2 & 3). This included positive detections from eight sampling sites, with at least one site being positive per water body (Table 2). Of the eight sampling sites that had positive detections for *S. ambigua*, six were likewise positive for *N. maculosus* (Table 2 & 3).

Within the metabarcoding data, two genetically distinct MOTUs were identified as *S. ambigua*. These two sequences have four bp differences over the 134 bp 16S amplicon region and display 3% genetic variation. The first, referred to as *Simpsonaias ambigua* 1, was detected only at the Wisconsin River sampling site, while the second, referred to as *Simpsonaias ambigua* 2, was detected in the other four watersheds (South Fork Licking, Licking River, Fish Creek, and Blanchard River) (Table 2). In addition to these genetically distinct female mitotype sequences, two genetically distinct male mitotype sequences were also detected for *S. ambigua*. These two distinct male mitotypes were also geographically split between the Wisconsin River and South Fork Licking/ Licking River. Similar to the female mitotype, these male mitotype sequences have four bp differences over the 138 bp 16S amplicon region and display 3% genetic variation. The complete metabarcoding dataset consisted of 1,591 sequence reads for the female mitotype of *S. ambigua*, whereas the male mitotype of *S. ambigua* comprised a total of 32,598 sequence reads (Table 2). Despite the apparent genetic variation across the sampled geographic range of *S. ambigua*, the qPCR assay targeting the mt-ND1 gene still provided positive detection for *S. ambigua* at locations for both mt-16S haplotypes. Metabarcoding and qPCR detections yielded similar results for estimating the presence of *S. ambigua*.

### Wisconsin River

The sampling at the Wisconsin River occurred at a site with the highest abundance of *S. ambigua* ever recorded in the state of Wisconsin, where hundreds of individuals were found beneath large sandstone slabs (Figure 2) (Stantec 2022). With eDNA, the female mitotype of *S. ambigua* was detected in nine of the 10 water samples with metabarcoding (Table 2). The sequence read abundance per water sample for the female mitotype ranged from 10 to 277, which accounted for less than 0.01% up to 0.06% of the total mussel DNA within a sample (Table 3). Whereas the male mitotype of *S. ambigua* was detected in eight of ten samples and was always detected at a greater sequence read abundance than the female mitotype, ranging from 18 to 7,740 reads per sample (Table 3). Additionally, *S. ambigua* was positively detected with qPCR in nine of the 10 water samples (Table 2). Two samples that were positive with metabarcoding for the female mitotype initially failed to yield a detection with qPCR and required an additional three technical replicates to obtain a positive detection with qPCR (Table 3). *Necturus maculosus* was likewise detected in nine of the 10 samples collected from the Wisconsin River site (Table 2).

**Figure 2.**
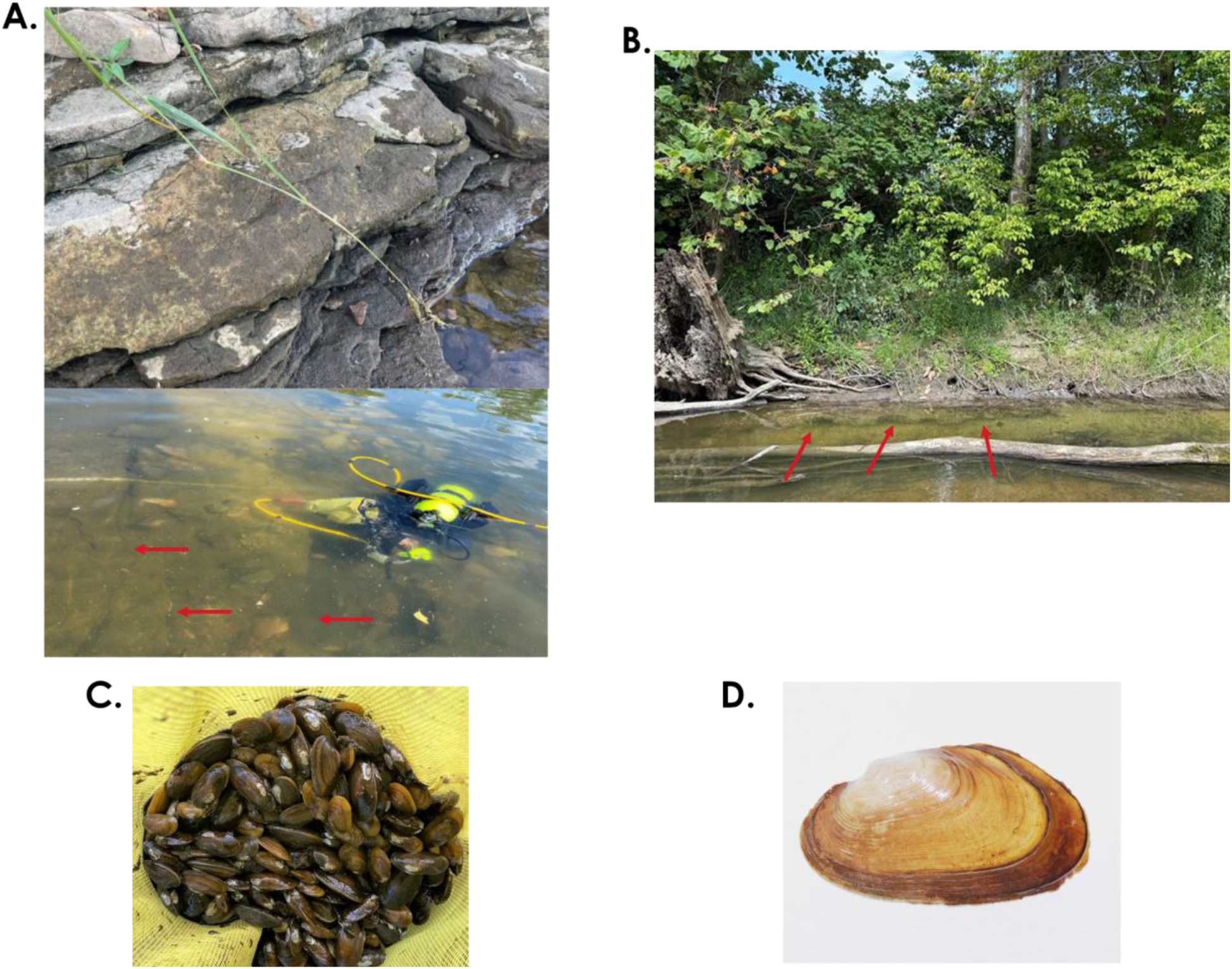
Observed habitat for Simpsonaias ambigua from (A) Wisconsin River where it occurs under sandstone slabs and boulders, and (B) Fish Creek where it occurs under overhanging American Sycamore (Platanus occidentalis) root clusters along the riverbank. Red arrows are used to illustrate the slabstone crevices or overhang of root clusters along the riverbank, respectively. Individuals collected from (C) Wisconsin River included the largest collection in the state of Wisconsin, and (D) Fish Creek where only two fresh dead shells were recovered. Photo credit: N. Berg (Stantec).

**Table 3.**
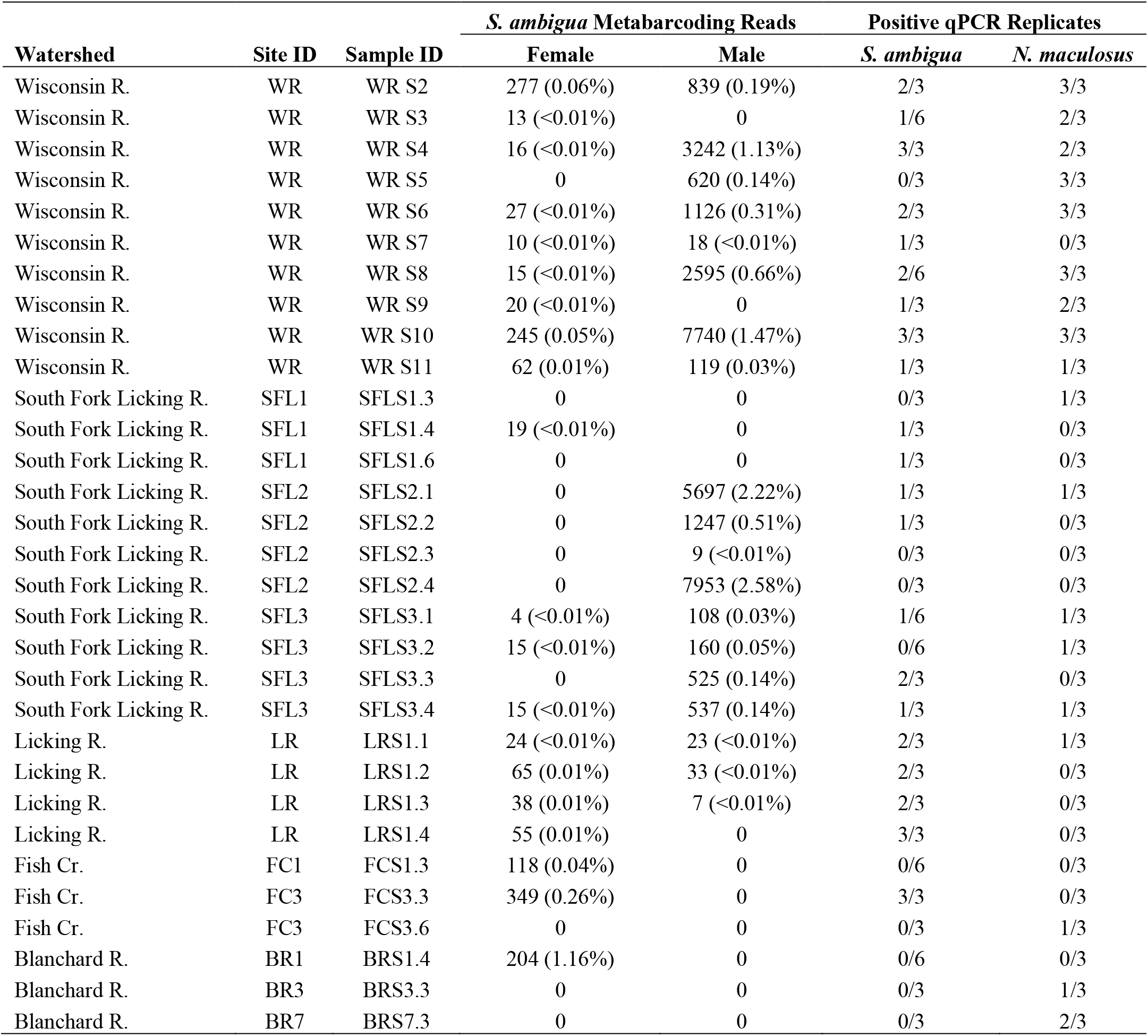
Environmental DNA detection of Simpsonaias ambigua using metabarcoding and qPCR methodologies and detection of Necturus maculosus using qPCR. Metabarcoding detection is listed as the sequence read count for both the female and male mitotypes. Parentheses indicate the percentage of the mussel DNA comprised of S. ambigua. qPCR detection is listed as the number of positive technical replicates out of the total tested. Samples with no detections for any method werex excluded from the table.

### South Fork Licking River

Visual surveys observed a single *S. ambigua* individual at both sites SFL1 and SFL2. These two individuals were recovered during the second pass of their corresponding search cell, and were found within adjacent search cells at the two sites (i.e., the last search cell of site SFL1 and the first search cell of SFL2).

With eDNA, *S. ambigua* was positively detected with metabarcoding in two of the three sampling sites, which included one positive sample at site SFLS1 and three positive samples at site SFLS3 (Table 2). The sequence read abundance of the female mitotype per water sample ranged from four to 19, which always accounted for less than 0.01% of the total mussel DNA within a sample (Table 3). The male mitotype of *S. ambigua* was detected in eight samples and from all three sampling sites, these detections included five samples that were non-detects for the female mitotype (Table 3). The male mitotype was always detected at a greater sequence read abundance than the female mitotype, ranging from nine to 5,697 reads (Table 3). Additionally, *S. ambigua* was positively detected with qPCR in seven of the 12 water samples, which included samples from all three sampling sites (Table 2). Two samples that were positive with metabarcoding for the female mitotype initially failed to yield a detection with qPCR. After the analysis of an additional three technical replicates, one of these samples provided a positive qPCR detection (Table 3). *Necturus maculosus* was detected in five of the 12 samples collected from the South Fork Licking River, which included at least one detection at each site (Table 2).

### Licking River

At the Licking River sampling site, *S. ambigua* was positively detected in all four samples with metabarcoding (Table 2). The sequence read abundance per water sample ranged from 24 to 65, which accounted for approximately 0.01% of the total mussel DNA within any sample (Table 3). The male mitotype of *S. ambigua* was detected in three of four samples and was always detected at a lower sequence read abundance than the female mitotype, ranging from seven to 33 reads (Table 3). With qPCR, *S. ambigua* was positively detected in all four samples. These qPCR detections all occurred in at least two of three technical replicates (Table 3). *Necturus maculosus* was detected in only a single sample collected from the Licking River (Table 3).

### Fish Creek

No live *S. ambigua* individuals were found during the visual surveys in Fish Creek. However, a focused tactile search of undercut banks beneath American Sycamore (*Platanus occidentalis*) root clusters uncovered two fresh dead shells (shells with indicators of recent death within the past few years) (Figure 2). These two fresh dead shells occurred just upstream of the FCS3 survey site. With eDNA, *S. ambigua* was detected with metabarcoding in two of the four sampling sites, which included one positive sample at site FCS1 and one positive sample at site FCS3 (Table 2). The sequence read abundance per water sample ranged from 118 to 349, which accounted for somewhere between 0.04% to 0.26% of the total mussel DNA within a sample (Table 3). Additionally, qPCR detected *S. ambigua* from FCS3 (Table 2). *Necturus maculosus* was detected in only a single sample collected from Fish Creek, which occurred at a site with detection of *S. ambigua* (Table 2).

### Blanchard River

Within the Blanchard River sampling sites, *S. ambigua* was not detected with visual tactile surveys or qPCR from any site. It was detected in one metabarcoding replicate but only for the female mitotype (Table 3). Detection occurred in only one of the 14 sampling sites, occurring at site BRS1 corresponding to most downstream location (Table 2). The sequence read abundance in this single detection was 204 and accounted for only 1.16% of the total mussel DNA within the sample (Table 3). *Necturus maculosus* was detected in one sample each from two of the 14 sampling sites in the Blanchard River, (Table 2).

### eDNA Detection Probability

Probability of eDNA detection estimates for *S. ambigua* were similar for qPCR and metabarcoding (Figure 3). The median estimated probability of detection for metabarcoding was 0.52 (0.35 – 0.69 CI), while that for qPCR was 0.61 (0.34 – 0.89 CI). Based on the confidence interval around the probability of detection, metabarcoding analysis would require the collection of three eDNA samples per site to exceed a 0.95 probability of detecting Salamander Mussel at the upper confidence to seven eDNA samples at the lower confidence. Similarly, to exceed a 0.95 probability of detection with qPCR analysis would require the collection of two eDNA samples per site at the upper confidence to eight samples per site at the lower confidence, plus the analysis of three technical replicates in the laboratory (Figure 3).

**Figure 3.**
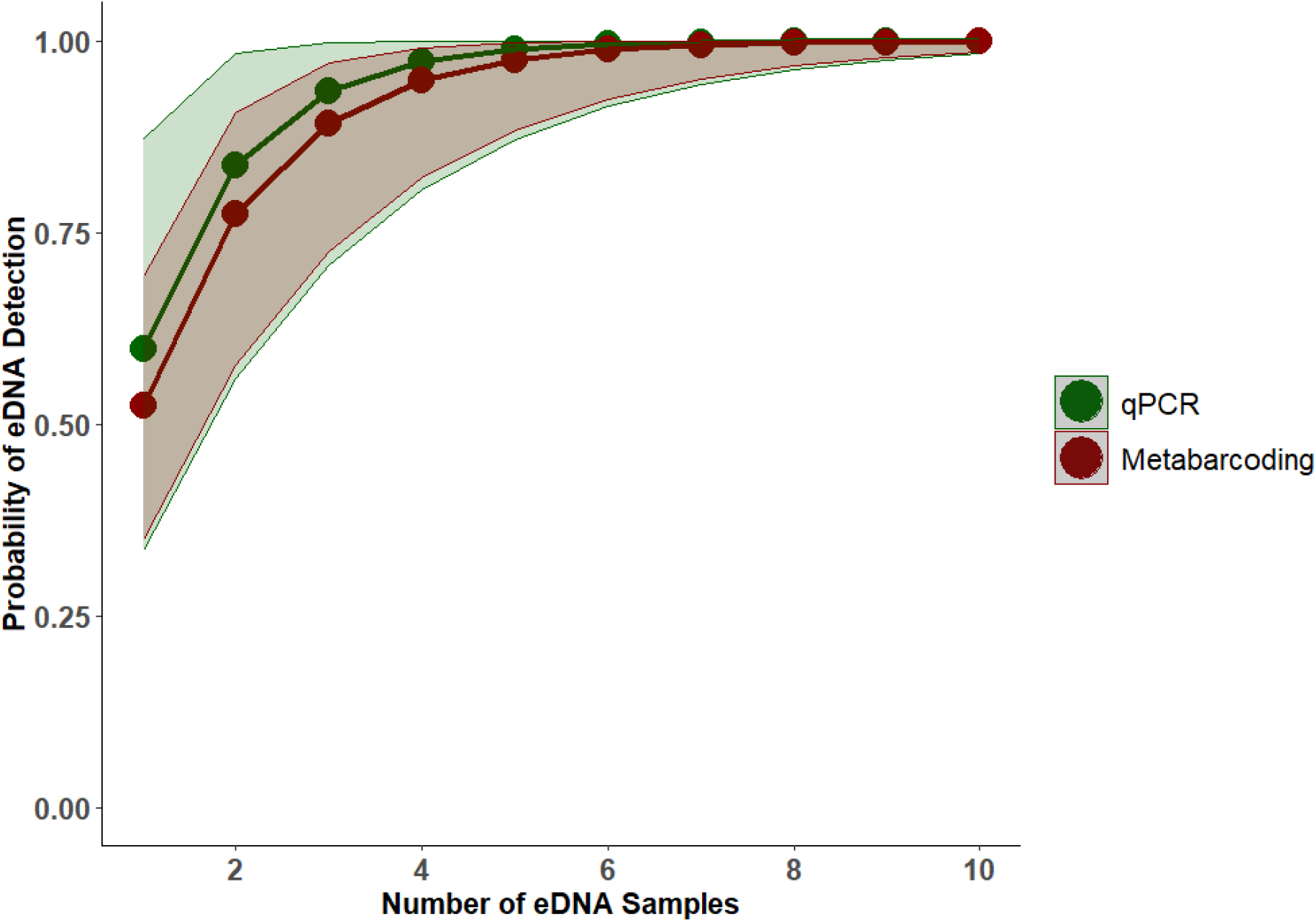
Environmental DNA probability of detection estimates for Simpsonaias ambigua when using metabarcoding or qPCR. Shaded region represents upper and lower confidence estimates. Estimates for qPCR are shown based on 3 technical replicates.

### Field and Laboratory Replication - qPCR

Both *S. ambigua* and *N. maculosus* were estimated to have a probability of detection exceed 0.95 when collecting at least eight eDNA samples per site and analyzing at least three technical replicates in the laboratory (Figure 4). Assessment of field and laboratory replication for qPCR analysis suggested collecting a greater number of samples in the field is more influential on increasing detection probability than analyzing more laboratory technical replicates per sample (Figure 4A & 4B).

**Figure 4.**
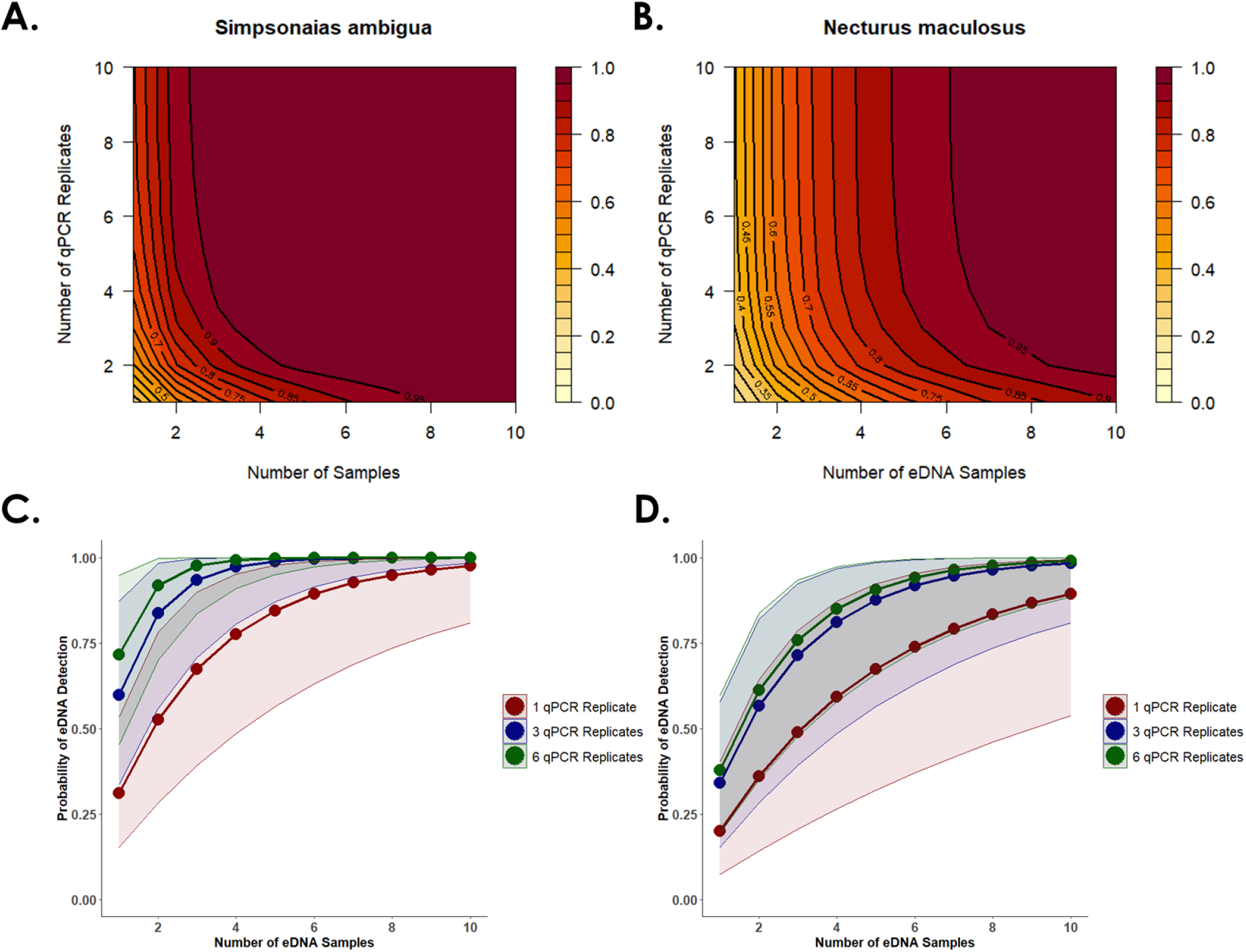
(A) The median estimate of probability of environmental DNA (eDNA) detection for Simpsonaias ambigua displayed across a range of survey effort and laboratory replication. (B) The median estimate of probability of eDNA detection for Necturus maculosus displayed across a range of survey effort and laboratory replication. (C) Probability of eDNA detection curves for S. ambigua. Shaded region represents upper and lower confidence estimates. (D) Probability of eDNA detection curves for N. maculosus. Shaded region represents upper and lower confidence estimates.

However, for both species, there was an initial probability of detection bump of nearly 2X when increasing the number of technical replicates from one to three (Figure 4C & 4D). For *S. ambigua*, the estimated probability of detection increased from 0.32 (0.15 – 0.54 CI) with one technical replicate to 0.61 (0.34 –0.89 CI) with three technical replicates. For *N. maculosus*, the estimated probability of detection increased from 0.20 (0.08 – 0.40 CI) with one technical replicate to 0.35 (0.17 – 0.57 CI) with three technical replicates. Yet increasing from three to six technical replicates yielded little additional detection power (Figure 4C & 4D).

## Discussion

*Simpsonaias ambigua* is a difficult species to detect using visual tactile survey methods, in part, due to specialized habitat use that differs from most other riverine unionids (i.e., they occur under boulders, within bedrock, and buried in the riverbank) (Porto-Hannes et al. 2025). Certain behaviors such as propensity for darkness, climbing ability, and clustering may compound detection difficulty. The current study produced positive detections from five sampling sites with recent visual confirmation of *S. ambigua* presence and across variable densities. These detections occurred in populations varying from a single individual in South Fork Licking River sites, up to a population of hundreds in the Wisconsin River (Stantec 2024). Additionally, detections occurred within Fish Creek from sites where visual surveys failed to recover live individuals with over 16 search hours. However, when an extra 10 hours of search was dedicated to digging through root wads within the riverbank, two fresh dead shells were recovered from a location of positive eDNA detection, demonstrating the sensitivity of eDNA.

### Detection efficiency and sampling effort

Contrary to these findings, previous eDNA surveys have reported relatively low detection rates for *S. ambigua* from sites of known occurrence in Ontario (Porto-Hannes et al. 2023). While overall detection rates were higher in the current study, the DNA signal for *S. ambigua* did appear to be low within all samples (measured as number of positive replicates in qPCR or the eDNA sequence reads in metabarcoding). Sporadic detection in replicates and low read counts are common for rare species (Marshall et. al. 2020). This was even true at the Wisconsin River sampling site, where samples were collected within a pocket of habitat that previously consisted of hundreds of individuals (Stantec 2024). It is worth mentioning that the previous survey in 2022 relocated 711 individuals as part of a mussel salvage effort, so the true size of this population at the time of eDNA sampling was unknown; however, it is still suspected to be large as the salvaged area was only a fraction of the available sandstone slab habitat present at the site (Fleece personal communication). For the majority of samples with a positive detection, the female mitotype of *S. ambigua* consisted of less than 0.01% of the total mussel DNA. Similarly, positive samples with qPCR rarely had detections occur in all tested technical replicates and were typically below the estimated limit of detection.

The apparent low concentration of *S. ambigua* eDNA may be due to their preferred habitat. For example, at the Wisconsin River site, Stantec (2022) found most of the *S. ambigua* between sedimentary sandstone layers, with clusters of *S. ambigua* sometimes observed in rock crevices 0.25 to 1.0 m away from open surface water. The degree to which surface waters mix with these interstitial waters is uncertain and could influence *S. ambigua* detectability. Additionally, previous laboratory studies have found comparable eDNA shedding rates between *S. ambigua* and other mussel species (Porto-Hannes et al. 2023), suggesting low eDNA concentrations at the Wisconsin River site are unlikely to be due to metabolic related reductions in eDNA shedding compared to other mussel species.

New technology and survey methodology requires proper evaluation of survey efficiency to estimate effort required to confidently assess presence / probable absence for a rare species (Marshall and Fleece 2025). Even though it appears that *S. ambigua* DNA was in low concentration within the environment, probability of detection estimates indicated relatively high detection probability can be reached with relatively low sampling effort (e.g., five to eight samples per site). This relatively low effort with eDNA is a fraction of the effort typically required with visual surveys. For example, the 26 hours of visual search in Fish Creek recovered only dead shells, yet eDNA detections were still achieved at two sites. The reduced field effort achieved through eDNA can be leveraged to sample a wider spatial extent, such as was possible with the spatial survey throughout the Blanchard River in which visual tactile surveys were conducted at only three sites while eDNA was collected from 14 sites. Overall, the lower field effort achieved with eDNA can lead to cost-efficiencies as projects scale in size (Bálint et al. 2018, Morris et al. 2024, Sternhagen et al. 2024)

As technology and equipment utilized for eDNA analysis rapidly expands, it is critical for proper communication of detection efficiencies for the implemented laboratory method (e.g., qPCR vs metabarcoding). Several studies have reported higher eDNA detection estimates for rare species when implementing species-specific assays using qPCR or ddPCR in comparison to community metabarcoding analysis, including assessments for invertebrates (Wood et al. 2019, Blackman et al. 2020) and vertebrates (Bylemans et al. 2019, Yu et al. 2022, Allen et al. 2023, McColl-Gausden et al. 2023, Kirtane et al. 2024). Species that occur at high DNA abundance within a sample can often mask the detection of rarer species during metabarcoding analysis, due to increased competition for limited PCR resources (Skelton et al. 2023). However, the more specific a metabarcoding assay is for its targeted taxonomic group, the better its ability to detect rare DNA targets (Marshall et al. 2020, 2024). The current analysis for *S. ambigua* found similar probability of detection estimates between the two laboratory methods.These results likely reflect the high specificity for unionid mussels targeted by the Prié et al. (2020) metabarcoding assay, which shows little to no cross-amplification for non-target taxa.

Seasonal fluctuations in environmental variables may influence eDNA detection rates (e.g., river discharge), with the current study occurring in late summer during low flows when dilution and transport effects are likely at their lowest (Curtis et al. 2021). Additionally, seasonal patterns in eDNA detection are often tied to the life-history and ecology of the target species (De Souza et al. 2016, Troth et al. 2021).Previous eDNA sampling for *S. ambigua* recovered higher detection rates in October than May (Porto-Hannes et al. 2023), although the overall sample size was low. The current survey occurred in August, potentially coinciding with timing of spawning for *S. ambigua* (Porto-Hannes et al. 2025), which may lead to heightened eDNA release.

### Presence or Probable Absence Assessments

Proper interpretation of eDNA results will always be more challenging than visual surveys, because you don’t have the species in hand to definitively conclude presence (Burian et al. 2021). In the current study, a total of 44 samples were collected from 13 sites throughout a 40.26 river km stretch of Blanchard River that has been proposed as critical habitat (USFWS 2023). This sampling effort in the Blanchard River yielded only a single positive detection for *S. ambigua* via metabarcoding analysis. This same eDNA sample did not provide a positive detection from qPCR across six technical replicates. This suggests the DNA signal was relatively weak within the sample in comparison to samples collected within the Wisconsin River, Licking River, and South Fork Licking River. However, this same detection pattern where a single positive detection occurred in metabarcoding and no detection occurred with qPCR was observed at Fish Creek, where the species was confirmed to be extant through the presence of fresh dead shells. Fish Creek and Blanchard River are both tributaries of the Maumee River watershed, where *S. ambigua* may embed within the riverbank.

A single positive detection of *S. ambigua* in contrast to dozens of negative samples in the Blanchard River requires due diligence before conclusive determinations can be made regarding presence or probable absence. A more intensive sampling effort focused on the single positive detection of *S. ambigua* at that location would be advisable to better understand the true presence of an extant population in this stretch of the river. Recent eDNA sampling in Illinois revealed an extant population of *S. ambigua* that hadn’t been recorded in over a century (Douglass et al. 2025), demonstrating the sensitivity of this technology. The data presented in earlier sections of this document suggest that a minimum of eight samples per site would be needed to have high confidence for detecting *S. ambigua*. Given the interest in conservation of the species plus the regulatory implications even more samples might be advisable.

While only a single detection occurred for *S. ambigua* within the Blanchard River, the detection of *N. maculosus* provides corroborating evidence for the possibility of an extant population. *Necturus maculosus* was detected in all watersheds and consistently detected at sites where *S. ambigua* occurred. Within the Blanchard River, *N. maculosus* was detected from two sites, including the Gilboa Quarry (site BRS3) which consisted of a long stretch of bedrock habitat, ideal for *N. maculosus* (Sutherland et al.2020, Porto-Hannes et al. 2025). In the current study, the eDNA detection probability of *N. maculosus* was estimated to be lower than that of *S. ambigua*, even though they are likely to reside within the same habitat. *Necturus maculosus* is most active during colder months, corresponding to the breeding season in late autumn and when they lay their eggs in early spring (Beattie et al. 2017). Considering the current eDNA survey occurred in late summer, the lower detection rates may reflect the reduced activity of *N. maculosus* during this time of the year. Previous visual surveys using setlines and minnow traps have noted higher capture rates during colder months for *N. maculosus* (McDaniel et al. 2009, Sutherland et al. 2020). Additionally, De Souza et al. (2016) recorded higher eDNA detection rates for *Necturus alabamensis* (Alabama Waterdog) in colder months than warm months. Regardless of sampling season, previous work in Michigan and Ohio have shown improved detection estimates of *N. maculosus* when using eDNA compared to traditional capture methods alone (Collins et al. 2019, Sutherland et al. 2020). The presence of host species for federally protected freshwater mussels is required to sustain a viable population, and thus the added value of eDNA to simultaneously assess presence of host and mussel will greatly aid in future conservation.

In addition to assessment of species presence, the metabarcoding analysis may be useful for analyzing metapopulation dynamics by evaluating genetic variation (Marshall et al. 2019). For example, the current study has identified two distinct sequences of *S. ambigua* across the geographic range of this study, with the eDNA detections in the Wisconsin River genetically differing from those detected elsewhere. The metabarcoding also produced differing sequences for the *S. ambigua* male mitotype between Wisconsin River and South Fork Licking River. eDNA sampling paired with metabarcoding can be important for identifying genetic variability and provide insight into distinct population segments that should be acknowledged in conservation planning (Parsons et al. 2025). Whereas qPCR analysis on its own does not provide the same genetic diversity information without follow-up sequencing of the PCR product (Goldberg et al. 2016). Additionally, the narrow specificity of qPCR can often lead to failed detection of a species when sampling across a large geographic range where unique genetic variants occur (Thalinger et al. 2021). Therefore, metabarcoding approaches for freshwater mussels can provide added value through the simultaneous detection of male lineages and genetic variation.

### Unionid male eDNA

Several studies have demonstrated large increases in eDNA concentration during spawning events for several non-unionid taxa (Tilliotson et al. 2018, Bayer et al. 2019, Thalinger et al. 2019), however it typically isn’t possible to differentiate between eDNA derived from somatic tissues versus gametes.Contrary, the unique DUI of the mt-genome in unionids makes it possible to draw inferences regarding eDNA signal associated directly from sperm release (see Marshall et al. 2025). Instead of analyzing the standard female lineage, detections of the male lineage (mostly restricted to sperm cells) may be required to document spawning behaviors for these taxa. By leveraging the metabarcoding assay in the current study, each sample could be simultaneously analyzed for both the female and male lineages (Marshall et al. 2025), whereas the employed qPCR assay was developed and validated for the female lineage alone (Porto-Hannes et al. 2023). In the current study, several samples yielded detections of the male lineage without detection of the female, and *vice versa*. Therefore, the metabarcoding approach applied here provides an added advantage of being able to detect either lineage if it is present in the sample.

In many cases, the detections of the male lineage for *S. ambigua* occurred in higher sequence abundance than that of the female lineage. This was especially true at the Wisconsin River site and two of the South Fork Licking sites, in which several samples had the male lineage occur at sequence read counts in the thousands compared to the female lineage in less than a hundred (Table 2). *Simpsonaias ambigua* is thought to spawn sometime in August thru October as females have been observed with developed eggs by August and gravid by September and October (Porto-Hannes et al. 2025). Therefore, the detections of the male lineage in early and late August may align with spawning. Interestingly, the initial survey site in South Fork Licking River (SFL1) occurred on August 19, 2025 and resulted in no detection of the male lineage for *S. ambigua*. However, the two subsequent survey sites (SFL2 and SFL3) sampled on August 20, 2025 and were heavily skewed towards the male lineage; yet these two sites were only 40 and 140 m upstream from SFL1, respectively. It is unclear why there was an apparent large increase in eDNA for the male lineage between sampling days, and further work is required to verify if this male signal truly originates from sperm presence.

## Conclusion

Environmental DNA surveying can be a powerful tool for the assessment of rare and cryptic species, both of which are often the case for *S. ambigua*. In the current study, eDNA detected *S. ambigua* from several sites of known occurrence. In the case of Fish Creek, no recent live individuals have been recovered, including during a recent multi-year survey effort (Marshall et al. 2026). Current eDNA detections led field personnel to subsequently devote greater search effort at precise locations to determine if previous detections could be substantiated. Through this increased effort, evidence of fresh dead shells buried within the riverbank was discovered. Without the initial molecular detections, it is likely this shell evidence of an extant population would have never been found. Detection probability estimates indicate populations of *S. ambigua* can be detected with high certainty using either metabarcoding or qPCR at relatively low sampling effort. This is an important finding, as metabarcoding is often considered worse for targeted detection of rare species. Furthermore, eDNA was used as an assessment of the population status throughout the Blanchard River, where there has never been a collection of any live individuals.The single detection in tandem with the presence of its host salamander, *N. maculosus*, indicates a population of *S. ambigua* within the Blanchard River may still be extant within the downstream portion of this unit, but further validation of this detection is required.

## Acknowledgments

This project was supported through data acquisition via the Kentucky Department of Fish and Wildlife Resources thanks to M. McGregor and the Ohio Department of Transportation (ODOT) thanks to M. Raymond, M. Michael, M. Perlik, and L. Scarberry

